# Open-source 3D printed air-jet for generating monodispersed alginate microhydrogels

**DOI:** 10.1101/804849

**Authors:** Dustin J. Hadley, Kevin T. Campbell, Marina H. Gabriel, Eduardo A. Silva

## Abstract

Open-source designs represent an attractive and new tool for research as it provides both affordable and accessible options to the lab environment. In particular, with the advent of new and cheap additive manufacturing technologies, the open-source design of lab hardware enables others to perform research that would be difficult otherwise. This manuscript describes an air-jet system designed to be open-source and simple to produce with 3D printing. The fully 3D printed air-jet was designed for the generation of hydrogel microbeads of a controllable size. Alginate microbeads were used as a working model, given that it has many promising research applications due to their injectability and highly reproducible properties. A fit definitive design of experiments was performed to determine critical factors affecting diameter, index of dispersity, and circularity of microbeads from this air-jet design. By regulating alginate concentration, air pressure, pump speed, and needle diameter could achieve control over microbeads size from 200-800 µm with low variance. Furthermore, we also demonstrate the potential probiotic research applications of the open-source air-jet through the encapsulation of bacteria in alginate microbeads with controllable degradation. The results of this study exhibit an open-source platform for making microscale biomaterials with controllable properties that can be achieved through budget 3D printers.

## Introduction

Publishing replicable scientific work remains recognized as a critical endeavor that researchers, funding agencies, and publishers should engage in cooperatively. Several efforts have been made to encourage reproducible science [4–6], including free open-source hardware (FOSH) and a growing number of online repositories, such as Github, Figshare, and Protocols.io [7, 8]. Specifically, FOSH cultivates participation in science by reducing supply limitations, fostering new research opportunities, and facilitating translation of these tools for educational purposes [8]. For example, open-source tools are currently available on the National Institutes of Health (NIH) website for use in cell cultures, microfluidics, and drug delivery systems [9]. Furthermore, the open-source license benefits researchers because building equipment yields a deeper understanding and allows modifications of designs [10]. The knowledge collected by users refines the open-source device to become robust while remaining accessible to others. One appealing strategy for creating FOSH is through designs that utilize 3D printing. The accessibility of fabrication with 3D printing has furthered the inventiveness within different communities, from laymen to aerospace engineers [10–15]. Moreover, the reduced cost of 3D printers in the past couple of years is resulting in 3D printers becoming available to the public at Makerspaces, public libraries, and universities [16].

One potential tool that would benefit from becoming a FOSH is an air-based system to produce microbeads. These microbead generators commonly employ polymer and crosslinker solutions in the formation of microscale hydrogel [17–20]. Microbeads are currently utilized and studied in different functions, including drug and cell delivery, cryopreservation, and scaffolds for tissue engineering strategies [20–22]. In these biomedical science applications, alginate is a particularly attractive polymer for microbeads, as it is biocompatible, undergoes rapid gelation under gentle conditions, and enables controllable mesh size [23]. Alginate, a copolymer comprised of (1,4)-linked β-D-mannuronate (M) and α-L-guluronate (G) residues, crosslinks in the presence of divalent cations, such as Ca^2+^, resulting in the formation of a 3D polymeric network characteristic of hydrogels [24–26]. Specifically, alginate microhydrogels have been applied to guiding the morphogenesis of progenitor endothelial cells and control the delivery of lentivectors and stem cells [27–29]. Alginate microbeads are commonly generated using droplet microfluidics, coaxial airflow units, two-channel air-jackets, and high voltage [22, 27, 28, 30].

To generate alginate microbeads, we designed a novel 3D printed air-jet system that generates control droplets of solution. An air-jet pushes air away from a source, making it similar to air bifurcation or electrostatic bead generators [22, 30, 31]. Our air-jet uniformly extrudes air across a needle attached to a syringe. A syringe pump supplied an alginate polymer solution at the needle tip, which formed droplets that fall into a calcium bath and generate hydrogel beads. Our objective for the device design was to permit modularity and sterility while having a low manufacturing cost by being compatible with budget 3D printers. We hypothesized controlling airflow, pump speed, and needle size for the 3D printed air-jet can create well-defined alginate microbeads. The device was characterized via a design of experiments (DOE) analysis using a fit definitive screening to define factors that altered resulting alginate bead characteristics. We expect this system to be reproducible between experiments and between laboratories due to well-defined set-up parameters that are typically less abundant in other publications using air-based droplet generating methods [31–33].

## Materials and Methods

### Device design and manufacturing

The air-jet was designed using Fusion 360® software (Autodesk, Inc) and prepared for 3D printing using Cura software (Ultimaker), an open-source slicer program. The device was split into three components, including the air-jet, the air intake, and supports. The air intake was printed with the air inlet parallel to the plane of the printing bed to prevent the nozzle from fracturing into the air tubing along with the printed layers. The air-jet was oriented with the outlet normal to the plane of the printing bed, such that the outlet for droplets was printed last, to prevent the need for support material. All components were printed on an MP Select Mini V2 (Monoprice, Inc) with poly (lactic acid) (Hatchbox®) at a layer thickness of 0.0875 mm and a printing speed of 50 mm/s. The extrusion temperature was 190 °C, and the heat bed temperature was 60 °C.

### Generation of alginate microbeads

LF 10/60 alginate polymer (~120-150 kDa) with higher G-block content (>60% as specified by the manufacturer) obtained from Novamatrix (FMC) was used to generate the microbeads. Alginate solutions were prepared by dissolving alginate polymer in either deionized water (diH_2_O) or phosphate buffer solution with added magnesium and calcium (PBS^++^; Life Technologies), as previously described [34]. The open-source air-jet and a syringe pump (Braintree Scientific) were run in parallel in a vertical position above a calcium bath. Syringes (Becton Dickinson) with varying needle sizes (PrecisionGlide™; Becton Dickinson) were positioned within the syringe pump and air-jet. The distance from the needle to the calcium bath was constant at 15 cm. The air-jet was then centered over the needle to generate a uniform air flow across the needle, with the tip of the needle slightly protruding from the outlet of air-jet. Nitrogen gas was then run through the air-jet before starting the syringe pump to prevent alginate from getting stuck in the air-jet chamber. After activating the syringe pump, the direction of droplets was checked using a flat surface. Adjustments were made in response to poor alignment of the needle through the center of the air-jet to ensure alginate droplets fell directly downwards. Finally, 5 mL of 100 mM calcium chloride (CaCl_2_) (Sigma) was used as the calcium bath for all experiments and placed underneath the air-jet. Alginate microbeads were generated for one minute with this setup before being analyzed directly from the wells using ImageJ software (NIH).

### Definitive Screen Design to characterize nitrogen pressure, needle gauge, and pump speed effect on microbead formation

A DOE approach with a definitive screening design (DSD; JMP software) was utilized to determine the impact of several factors on the properties of microbead generated with this air-jet design. A fit definitive screening test was used to ascertain active main effects and second-order effects of buffer type, needle diameter, air pressure, alginate concentration, and pump speed on bead diameter, the index of dispersion of bead diameters, and circularity. The effect of polymer solution viscosity was investigated using alginate dissolved at different concentrations (1, 2, and 3% (w/v)) and in different solvents (PBS^++^ and diH_2_O). The various needle gauges of 27, 21, and 18 were tested, which had approximate diameters of 0.210, 0.524, and 0.840 mm, respectively. Nitrogen pressure was altered between 200 to 800 kPa using a pressure gauge attached to a compressed nitrogen gas tank. Syringe pump speeds were also varied between 100, 250, and 400 µL/min. Index of dispersion is the variance normalized to the mean and was used to analyze bead uniformity. The circularity is defined below in the following equation:

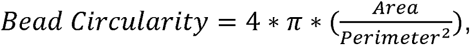

where the value of bead circularity varies between 0 to 1, with 1 being a perfect circle. Microbeads were analyzed with ImageJ, and significance was determined via the fit definitive screening analysis. DSD utilized a minimum of 18 groups, where a group is a set of differing factors, with n = 8 samples per group. The residual was calculated for each examined factor by subtracting the mean of all measured values of a specific factor from the average of a specific condition of the factor. If conditions of a factor are determined to be significantly different, trends were described with predictive plots generated using least-square regression or polynomial least squares.

### Additional experiments on experimental design parameters

Additional experiments were performed to determine how the influence of air pressure and pump speed on microbead formation. Air pressures were used from 0 to 800 kPa in 100 kPa increments. For 0 kPa, the air-jet was removed to prevent alginate solution from getting stuck within the device. The pump speeds of 10, 50, 100, 150, and 200 µL/min were used with a calcium bath. The viscosity and surface tension of the calcium bath was altered through the addition of surfactant, TWEEN® 20 (0.01% (v/v)), in pump speed experiments. Both experimental set-ups utilized a 27G needle with 2% (w/v) alginate, and 100 mM CaCl_2_ baths. Bead diameter, diameter index of dispersion, and circularity were determined using ImageJ (NIH).

### Bacteria in alginate microbeads and alginate lyase bead degradation

Green fluorescent protein (GFP) expressing *Escherichia coli (E. coli)* were loaded into 2% (w/v) alginate solution and dropping into 100 mM CaCl_2_ solution containing 0.25% (w/v) chitosan (Sigma). Chitosan was initially dissolved at 4% within 0.066 M glacial acetic acid (Sigma) before being diluted to the final concentration. Microbeads were made by transferring alginate solution through a 25G needle at 100 µL/min with the air-jet fed by nitrogen gas at 600kPa. *E. coli* loaded microbeads were cultures in Lysogeny broth (LB; VWR). For the generation of degradable alginate microbeads, alginate polymer was incorporated with alginate lyase, an enzyme that cleaves glycosidic bonds in alginate polymers [35], at various concentrations (0, 5, and 50 mU/mL). A syringe with a 27G needle was filled with an alginate solution, containing alginate lyase, and microbeads were generated with the air-jet as described above. These alginate microbeads were left to gel for 15 minutes and subsequently washed in DI water. Next, approximately 100 µL of beads were transferred to 12-well plates, topped in 4 mL of EGM-2MV media (Lonza), and incubated at 37 °C. At 1 hour and 1, 3, 5, 7, 14, and 21 days microbeads were imaged, and the size of the beads was analyzed with ImageJ. The initial distribution of microbead sizes at 1 hour, and the change in microbead size over 21 days was reported (n = 50).

### Statistical analysis

Comparisons were assessed by Student’s unpaired t-tests. Differences between conditions were considered significant if P < 0.05. All analyses were performed using GraphPad Prism software (GraphPad Software) and JMP software.

## Results

### Design of a 3D printed air-jet

The air-jet system was engineered for the rapid generation of alginate microbeads with consistent and controllable size. The design of the air-jet was optimized for a Fused Deposition Modeling (FDM) 3D printer, which enabled the internal features to be printed without support material. The distance between the inner cylinder that holds the needle and the outer wall was 1 mm to prevent fusing between features while printing (Fig 1A). Additionally, air flows around this gap (teal), which has a cross-sectional area of approximately 33 mm^2^. This area expands to a maximum of 103.9 mm^2^ before being constricted to an area of 7.1 mm^2^ at the exit hole (green). The small needle channel (orange) has a cross-sectional area of approximately 1.77 mm^2^, which is further reduced with the inserted needle. This region reduces air from traversing up the needle inlet and increases air velocity at the tip of the needle. Including an angled inlet (red), the overall profile of the air-jet is compact to allow needles to fit through the air-jet (Fig 1B). Furthermore, the construction permits the incorporation of a needle and syringe, either through direct attachment to the syringe pump or separately through adjustable stands (Fig 1C-D).

**Fig 1.**
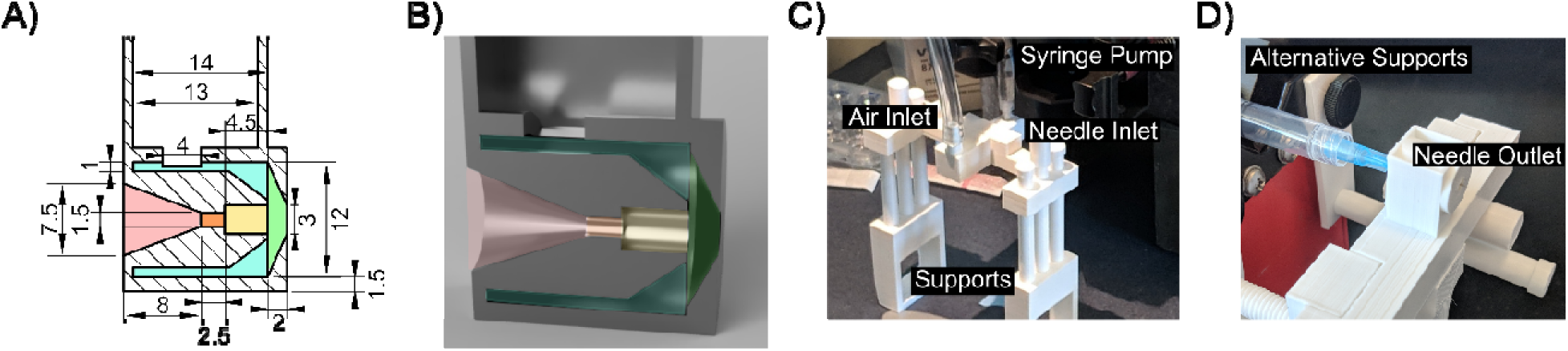
3D Printed Air-jet design and set-up. An engineering drawing of the internal dimensions displays an opening for airflow and the needle (A). A rendering of the model cross-section (left) and external view (right) (B). The 3D printed air-jet can be manipulated into different positions via the adjustable leg supports (C) or held in place by attaching directly to the syringe pump (D).

### Fit Definitive Screening Test

A fit definitive screening test was performed to determine the main active effects of buffer type, needle diameter, air pressure, alginate concentration and pump speed on bead size, the variance of bead sizes, and circularity. Residuals for these factors were calculated and statistically assessed for first-order or second-order correlations. (Figure A-C in S1 Figure). From this definitive screening design, prediction trends were calculated for factors exhibiting a significant first-order or second-order relationship with microbead size, variance, or circularity. The equation for these predictive trends of continuous factors are presented in each subfigure (Fig 2A-C). Alginate concentration was determined to have a second-order effect on bead diameter, with higher alginate concentrations resulting in increased bead diameter (Fig 2A), decreased bead variance (Fig 2B), and increased circularity (Fig 2C). Similar trends were also observed with a decrease in air pressure. Needle diameter had a small effect on bead diameter, with a smaller needle diameter creating smaller beads. Additionally, the buffers used to dissolve the alginate polymer were found to influence the index of dispersity, with PBS^++^ usually leading to a larger distribution of bead sizes. Representative images of the beads under similar conditions are shown. Beads with lower alginate concentrations were smaller and less uniformly round (Fig 2D). Using a faster pump speed, from 250 to 400 µL/min, led to more uniformly round beads that appear slightly larger (Fig 2E). Beads formed with alginate dissolved in diH_2_O appear more uniformly round compared to alginate dissolved in a phosphate buffer (Fig 2F), while beads were smaller using a smaller needle size (Fig 2G).

**Fig 2.**
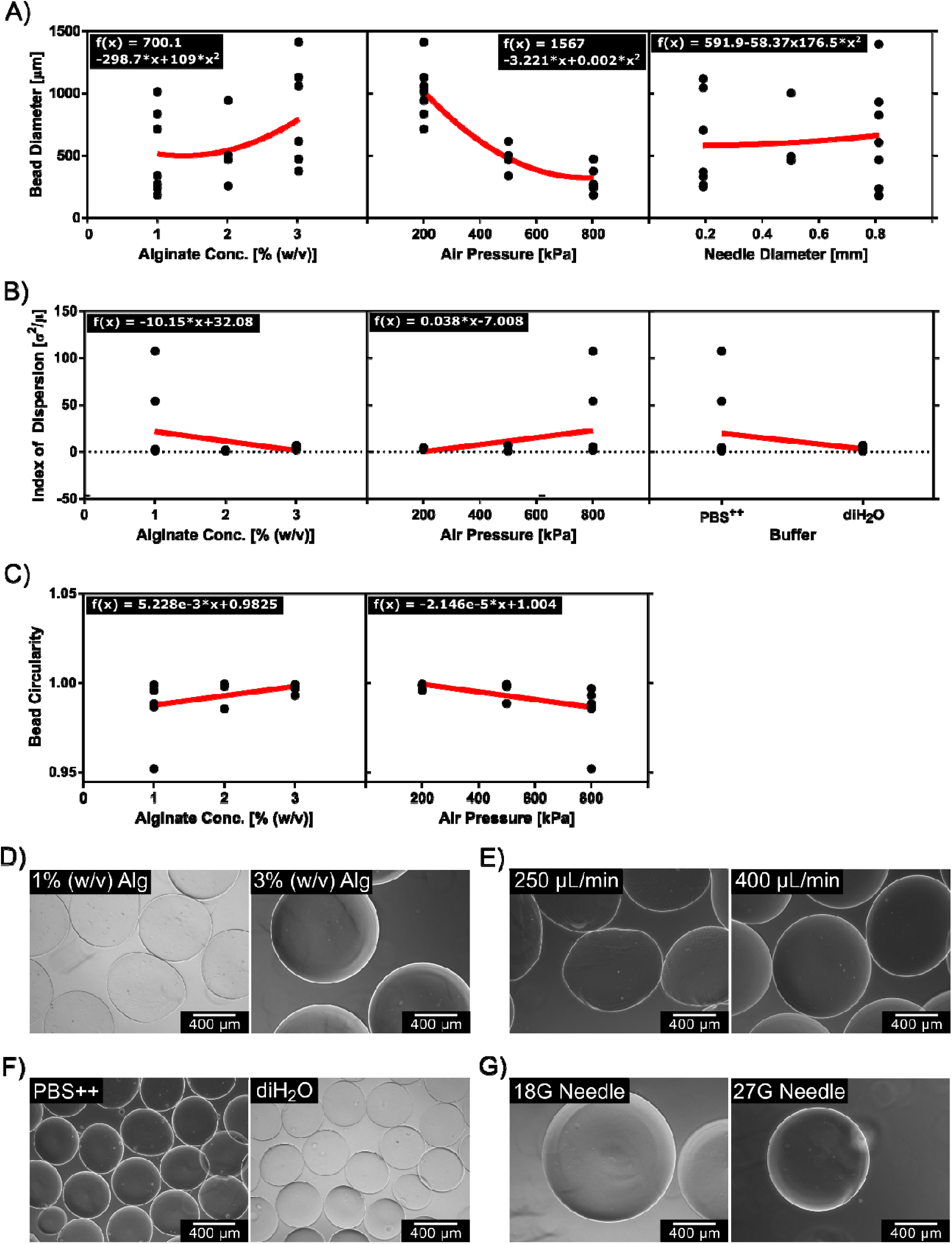
Prediction plots of factors affecting alginate microbead formation and representative images. Prediction values combine groups with a specific factor condition regardless of differences in other factor conditions. Red lines represent linear or nonlinear fits, determined by the statistical significance of first-order or second-order effects. The outcomes of interest were bead diameter (A), index of dispersion of bead diameters (B), and circularity (C). Comparable groups of microbeads generated from the design of experimental groups were displayed. A 27G (0.21 mm) needle with 200 kPa of air was used to dispense alginate at 1 and 3% (w/v) at a rate of 400 µL/min (D). Lastly, an 18G (0.838 mm) needle was used to dispense 1% (w/v) alginate at 250 or 400 µL/min using 200 kPa of air (E). Different buffers were tested with a 21G (0.534 mm) needle supplied with 2% (w/v) alginate at 250 µL/min with 500 kPa of air flowing over the needle (F). Next, 18G (0.838 mm) and 27G (0.21 mm) needles were tested with 100 µL/min of 3% (w/v) alginate with 200 kPa of airflow (G). In A-C, points represent the mean value, from n = 8 samples, of individual runs of a design of experiments.

### Characterization of Microbeads from changes in Air Pressure and Pump Speed

Next, we further investigated air pressure and pump speed as possible factors for controlling bead sizes. A 27G (0.21mm) needle was used to make beads with 2% (w/v) alginate. The addition of airflow had a drastic effect on the bead size (Fig 3A). From the initial addition of airflow, the bead size decreased with a one-phase decay (R^2^ of 0.9925). The equation for the nonlinear fit is

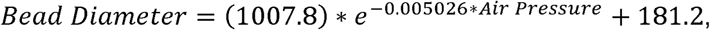

where air pressure is in kPa. For the range where airflow was used, the decrease in bead size with increasing air pressure is approximately linear (R^2^ = 0.8165), described by the following equation:

**Fig 3.**
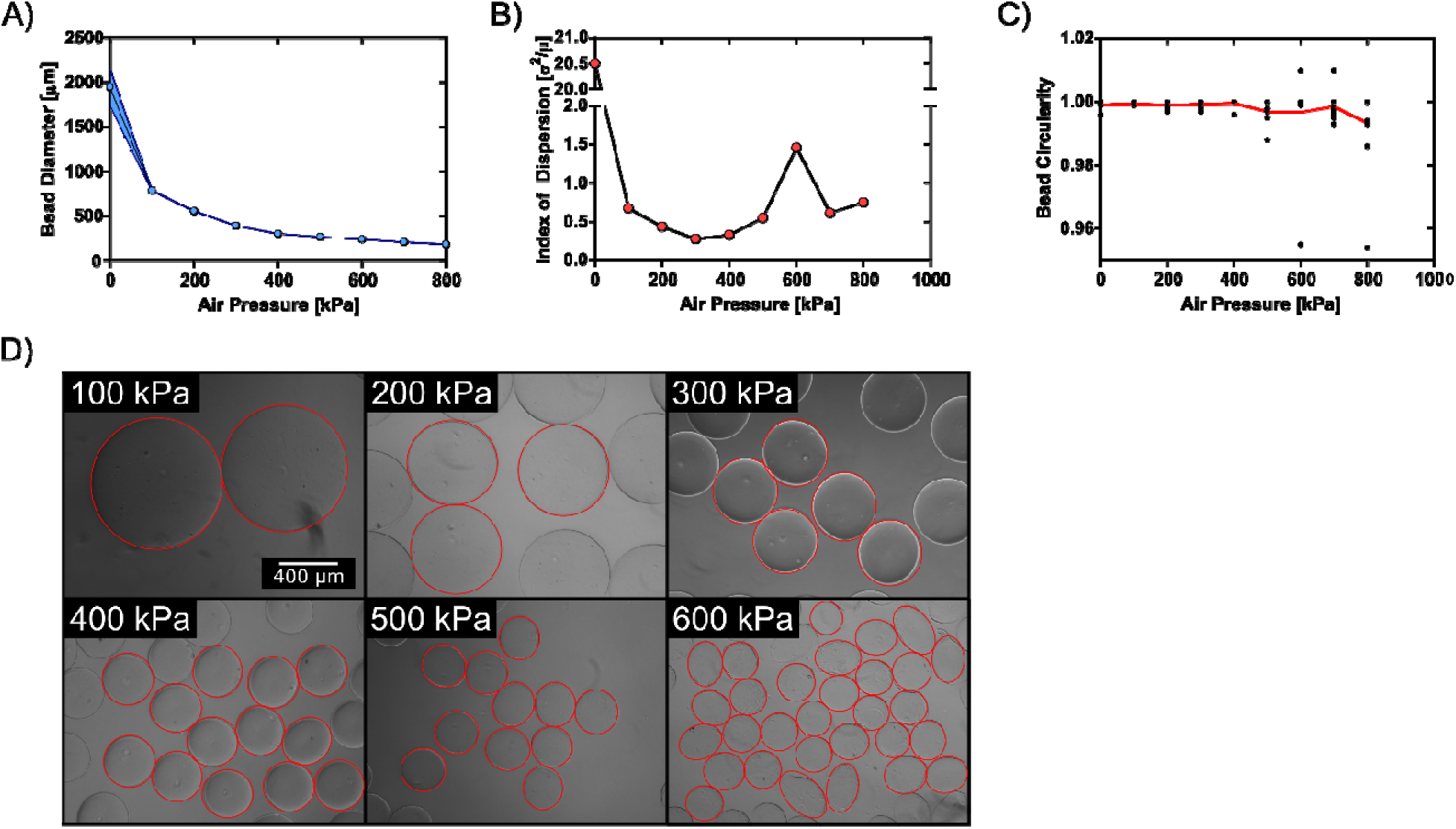
Characterization of air pressure effect on alginate microbeads. Beads were generated using a 27G needle (0.21 mm diameter) with 2 % (w/v) alginate 10 cm from the calcium bath. The microbeads were generated with changing air pressure (A-C) with a constant pump speed (100 µL/min). Generally, an increase in air pressure led to a decrease in the microbead diameters (A) and circularity (C). From the microbead diameter samples, the index of dispersion was calculated for bead diameters (B). In figure panel A, the mean is represented by the central line with the shaded areas denoting the error envelope of the standard deviation (n = 9). In figure panel B, points represent the index of dispersion of microbead diameters, calculated with the mean and standard deviation from the samples in panel A. In figure panel C, each data point represents the circularity of an individual microbead, that were also measured for their diameter (n = 9), with the mean of each condition represented by the central lines. Representative images of microbeads from air pressures 100-600 kPa are displayed with red outlined to highlight the resulting change in diameter and uniformity (D).

Increasing nitrogen gas pressure was found to both decrease bead size and increase the index of dispersion (Fig 3A-B). While no airflow creates beads with the largest index of dispersion, the variance in bead diameters increased as air pressure is increased beyond 400 kPa (Fig 3B). A rise in air pressure beyond 400 kPa also led to a reduction in bead circularity and greater deviation in circularity (Fig 3C). Representative images of these beads show drastic changes in bead diameters with the addition of air and loss of circularity at higher air pressures (Fig 3D).

The addition of surfactant to the calcium bath was also determined to effect microbead properties. Alginate microbeads were created with various pump speeds using a 2% w/v alginate polymer solution and 400 kPa air pressure. The bead diameters were determined to linearly decrease with pump speed, R^2^ = 0.8427 (Fig 4A). The bead diameter increased by approximately 40 µm per 100 µL/min change in pump speed in the range tested regardless of whether a surfactant was used or not. There is no significant difference in the index of dispersion of bead diameters with the addition of surfactant, nor for bead circularity (Fig 4B-C). Together, these data demonstrate how bead diameter formation is dependent on factors that occur at the air-jet, while circularity of alginate microbeads is adjusted through factors relating to the calcium bath.

**Fig 4.**
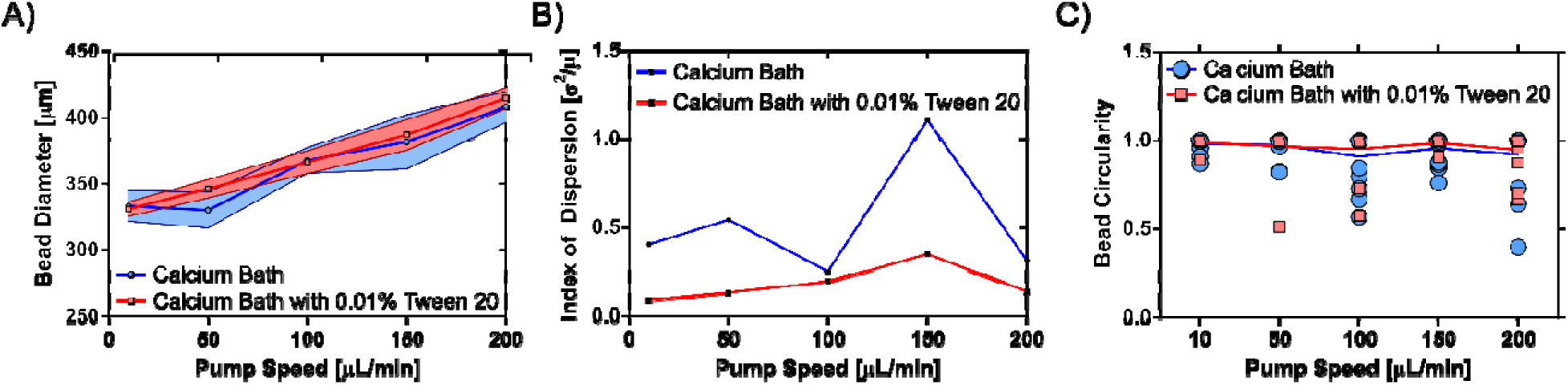
Effect of pump speed and calcium bath surface tension on microbeads formed via the 3D printed air-jet. The diameter of microbeads increased with quicker pump speed, at a constant air pressure, regardless of whether the calcium bath included a surfactant (0.01% (v/v) Tween 20)) (A). Furthermore, microbead uniformity was slightly improved through the addition of the surfactant, as indicated by the consistently lower index of dispersion and circularity. In figure panel A, the mean is represented by the central line with the shaded areas denoting the error envelope of the standard deviation (n = 6). In figure panel B, points represent the index of dispersion of microbead diameters, calculated with the mean and standard deviation from the samples in panel A. In figure panel C, each data point represents the circularity of an individual microbead, that were also measured for their diameter (n = 6), with the mean of each condition represented by the central lines.

### Validation of Open-Source Air-jet System Utility for Encapsulation Strategies

Alginate microbeads generated from the air-jet was interrogated for potential biomedical applications. Bacteria were successfully encapsulated in alginate microbeads of varying diameters by changing the air pressure (Fig 5A). Furthermore, these bacteria were viable and could proliferate within the alginate microbeads (Fig 5B). The air-jet could also generate degradable alginate microbeads. Alginate microbeads containing various alginate lyase concentrations were found to initially have similar distributions of bead diameters (Fig 5C). The alginate microbeads containing alginate lyase decrease in size over 21 days, with the largest change within the first few days (Fig 5D). An approximate 27.3% reduction in microbead diameter was observed with 50 mU/mL of the enzyme.

**Fig 5.**
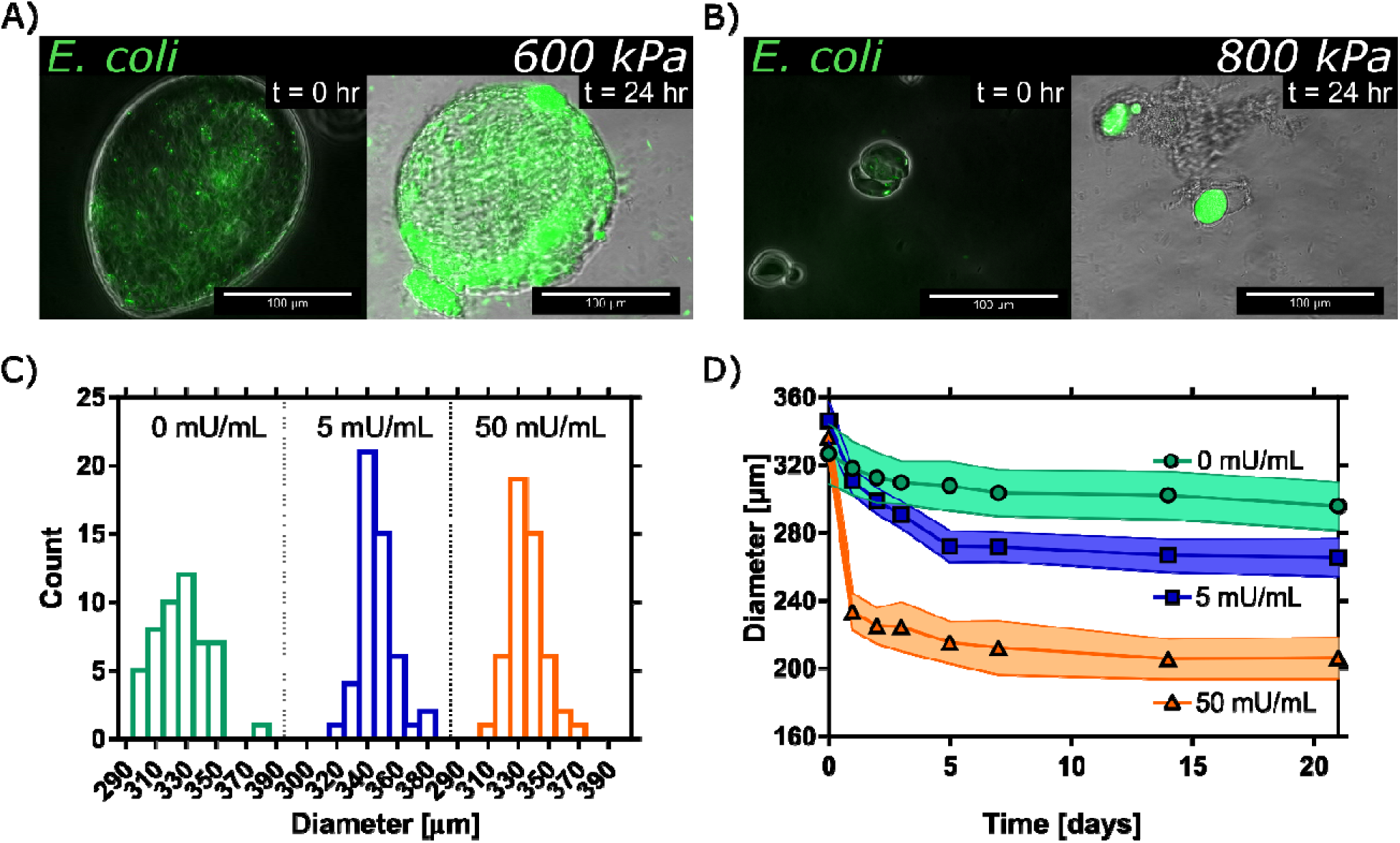
Encapsulation of bacteria and alginate degrading enzyme. GFP expressing *E. coli* were encapsulated in alginate microbeads through applying the air-jet with varying air pressures, 600kPa (A) and 800 kPa (B), and imaged for the change in fluorescence between 24 hours. In both cases, the enhanced distribution of the bacteria is observed over this timespan. In addition, varying concentrations of alginate lyase were encapsulated in alginate microbeads utilizing the air-jet system. The diameters of 50 microbeads per alginate lyase concentration suggest no initial difference in microbead size due to the encapsulation of the enzyme (C). Enzymatic degradation leads to a reduction in microbead diameters (D). In figure panel D, the mean is represented by the central line, with the shaded areas denoting the error envelope of the standard deviation (n = 50).

## Discussion

The designed open-source air-jet reliably generated microbeads without the need for complex assembly or expensive lab equipment. To our knowledge, this is the first fully 3D printable air-jet system. Importantly, this device controls needle placement and airflow, making set-up easy and reproducible compared to other air-jet systems [18, 36]. Furthermore, the geometries of internal features take advantage of FDM 3D printers to generate the air-jet as one piece benefiting the accessibility and affordability of the design. Overall, the described air-jet system offers control over microbeads with defined set-up parameters that can be created with budget 3D printers.

The open-source air-jet described here provides an advantage in ensure reproducibility and reliability by allowing defined set-up parameters. Specifically, the separation from the needle and the airflow, comparable to other air-jets that shear droplets off the needle tip; however, the fixed needle position ensures central alignment with the airflow [20, 33, 37]. Consequently, the described air-jet requires consideration of needle size is important to prevent the backflow of solution or air. In contrast, electrostatic bead generators do not need to consider needle size or backflow of components, but the air-jet is still advantageous for not requiring an electrical power source [18, 38]. To further prevent possible backflow or entrapment of solution, smaller diameter holes at the needle inlet were designed. Additionally, the air was flowing through the device before running the syringe pump with the needle placed approximately 0.5 mm from the outlet. Together, these considerations result in monodispersed alginate microbeads in minutes, even with highly viscous solutions, for what would take hours with microfluidics [28, 39].

The fit definitive screening test determined polymer concentration, air pressure, needle diameter, and buffer type as factors influencing microbead formation. Advantageously, the DOE methodology dramatically reduced the total number of groups, from 162 to 18, where a group represents a specific combination of different factor values that could manipulate microbead properties. The results suggest that a higher alginate concentration, larger needle diameter, or lower air pressure leads to larger circular beads, an outcome consistent with alginate microbeads formed via other air-bifurcation or air-jet systems [17, 18, 37, 39]. Whereas lower concentrations of alginate and higher airflow speeds could generate smaller beads, the prediction trends suggest this could lead to problematic increase variance and decrease circularity. Although the circularity was improved with higher alginate concentrations, higher polymer densities can be disadvantageous for cell applications where cells are required to migrate through the hydrogel scaffold [37, 39]. Therefore, alternative methods, such as air pressure and needle size, should be utilized with the optimal polymer concentrations to achieve circular hydrogel microbeads of the desired diameter. Taken together, alginate microbead properties can be controlled by adjusting multiple parameters involved with this air-jet design.

Pump speed and air pressure were interrogated as methods to regulate microbead size, circularity, and uniformity. While air pressure could control the size of beads from 2 mm to 200 µm, adjusting pump speed changed bead sizes from 350 to 425 µm, suggesting air pressure provides a more dynamic range of bead sizes. However, both methods generate beads in the range of 200-800 microns in diameter with a coefficient of variation of less than 10% that is often desired for encapsulation of cell applications [40]. Although microbeads smaller than 200 microns can be made with higher air pressures, tear-shaped droplets will form without sufficient distance and time for the droplet to become round [41]. Additionally, the surface tension of the bath also contributes to deviations in circularity. Consistent with previous reports, the addition of a surfactant to the calcium bath did improve the circularity of beads in this system [39]. Overall, the designed open-source system generates microbeads with desirable characteristics, with similar considerations as other air-jet systems being required to improve bead uniformity.

The encapsulation of bacteria and the enzyme alginate lyase highlight the distinct benefits of the described air-jet system for encapsulating cargo that can degrade the hydrogel matrix. Here, we load microbeads with bacteria, and their growth over 24 hours results in pockets of bacteria within the hydrogel matrix. The encapsulation of bacteria is promising for numerous biomedical applications, including probiotic delivery, especially for anaerobic bacteria sensitive to atmospheric oxygen concentrations [21, 42–45]. By using the air-jet system with nitrogen gas in a nitrogen-enriched chamber, microbeads could be rapidly generated with anaerobic bacteria while displacing oxygen [21]. Furthermore, the use of enzymes to degrade alginate has been previously used to control the delivery of endothelial progenitor cells and adeno-associated vectors [46, 47]. The rapid degradation of microbeads observed here suggests heterogenous microbeads could result from a slower output rate, such as with microfluidic methods [28]. Herein, we demonstrate the specialized utility of this open-source air-jet system that can rapidly generate monodispersed microbeads for encapsulation applications.

In conclusion, we present an open-source air-jet with simple features that can be printed on a small budget 3D printer. The system’s modular design allows easy cleaning, interchangeability to customized set-ups, and rapid replacement. We have demonstrated that alginate microbeads can be generated using this open-source air jet system with reasonable microbead diameters and variance. We anticipate the system will enable other research laboratories, as well as other fields, to easily generate polymeric microbeads.

## Supporting information

Supplemental Figure 1 - Residual Plots

Supplemental Table - Bill of Materials

## Acknowledgments

Funding for these studies was provided by the American Hearth Association (17IRG33420114)

