## Supplemental Figure 1 - Residual Plots for "Open-source 3D printed air-jet for generating monodispersed alginate microhydrogels"

**S1 Fig. DOE residual plots for determining key factors on alginate bead characteristics.**

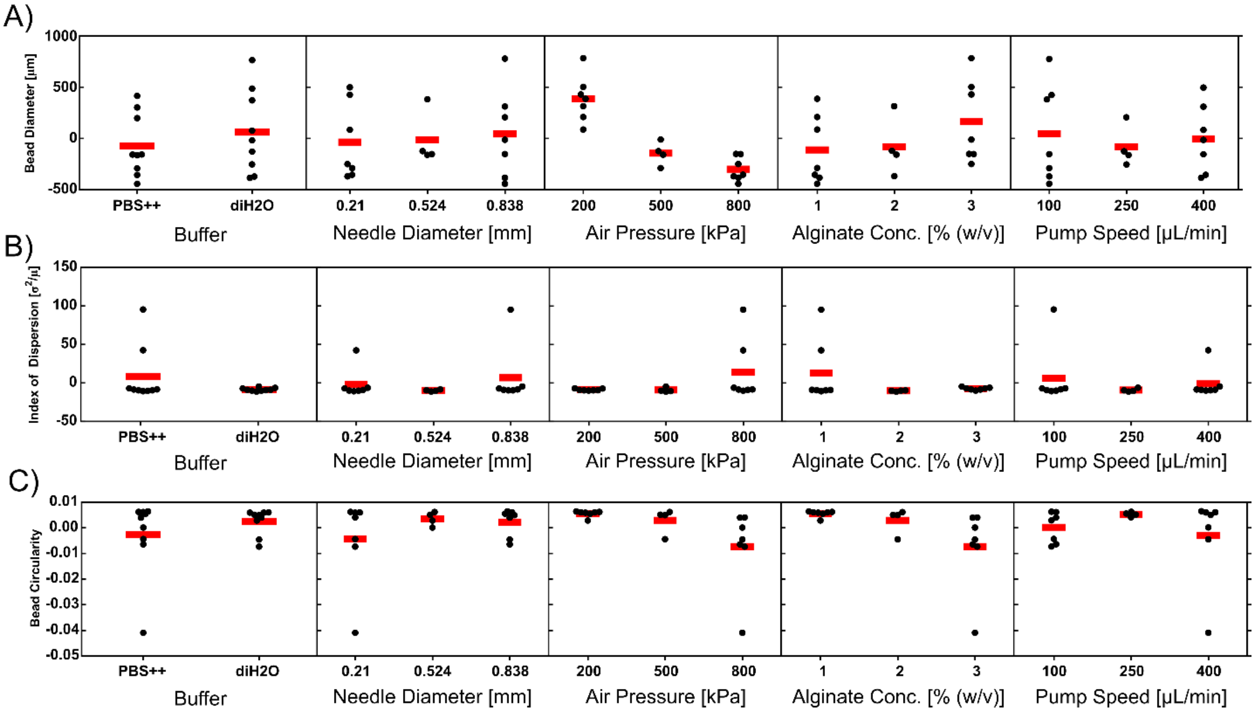


A fit definitive screening test was performed to determine important factors for bead diameter (A), index of dispersity for bead diameter (B), and bead circularity (C) outcomes. Factors tested include solute of phosphate buffer ($PBS^{++}$) and $diH_2O$, needle diameter, air pressure, alginate concentration, pump speed. Eighteen runs, representing unique combinations of the factor values, were performed with n = 8 samples per run. The samples were averaged to represent the mean value of the specific run condition. Residuals were then calculated from the subtraction of this mean value from the mean of all values for that factor (shown as black dots); i.e., air pressure or pump speed. The red bar represents the mean residual value for each conditional value. The deviation of these means from zero suggests factors that influence the outcome.
