## Supplemental Table - Bill of Materials for "Open-source 3D printed air-jet for generating monodispersed alginate microhydrogels"

### S2 Table. Bill of Materials

| Item | Details | Price | Alternatives |
|---|---|---|---|
| *FDM 3D Printer* | Monoprice Mini V2 | $220 | Any 3D printer technology should be compatible. Public access to 3D printers exists to make this not a necessary expenditure. |
| *PLA filament* | Hatchbox PLA filament, white, 1.75 mm, 1kg | $20 | Additional brands or material types can be used. |
| *Syringe pump* | Just Infusion Syringe Pump | $275 | Open-source syringe pumps have been previously described. |
| *Syringe needles* | 18G (100 pack) | $14 | Different sizes are compatible. |
| *Syringe* | 5 mL with luer lock (100 pack) | $14 | Any size capacity is compatible. |
| *Beaker* | Used for collecting microbeads, 50, 100, and 250 mL pack | $9 | Any liquid container works. Alternatively, we have used 50 m conical tubes. |
| *Nitrogen gas* | | $40 | House air supply or any pressurized air can be used. |
| *Gas regulator* | Nitrogen gas regulator | $60 | |
| *Tubing* | Used for gas lines | $18 | |
| *Hose Clamp* | Used to tighten tubing on gas regulator (20 pack) | $8 | |
